# “Stick-slip dynamics of cell adhesion triggers spontaneous symmetry breaking and directional migration”

**DOI:** 10.1101/354696

**Authors:** K. Hennig, I. Wang, P. Moreau, L. Valon, S. DeBeco, M. Coppey, Y. A. Miroshnikova, C. Albiges-Rizo, C. Favard, R. Voituriez, M. Balland

## Abstract

Directional cell motility during organism and tissue development, homeostasis and disease requires symmetry breaking. This process relies on the ability of single cells to establish a front-rear polarity, and can occur in absence of external cues. The initiation of migration has been attributed to the spontaneous polarization of cytoskeleton components, while the spatiotemporal evolution of cytoskeletal forces arising from continuous mechanical cell-substrate interaction has yet to be resolved. Here, we establish a one-dimensional microfabricated migration assay that mimics complex *in vivo* fibrillar environment while being compatible with high-resolution force measurements, quantitative microscopy, and optogenetics. Quantification of morphometric and mechanical parameters reveals a generic stick-slip behavior initiated by contractility-dependent stochastic detachment of adhesive contacts at one side of the cell, which is sufficient to drive directional cell motility in absence of pre-established cytoskeleton polarity or morphogen gradients. A theoretical model validates the crucial role of adhesion dynamics during spontaneous symmetry breaking, proposing that the examined phenomenon can emerge independently of a complex self-polarizing system.

**One sentence summary:** Cells can autonomously break their symmetry through traction force oscillations (mechanical instabilities) that lead to stochastic detachment of adhesion patches on one side of the cell and the subsequent initiation of migration.

## Main text

Directional motility is a plastic process (1) that is the fundamental basis of key biological processes in eukaryotes, such as embryonic morphogenesis, leukocyte trafficking in immune surveillance, and tissue regeneration and repair (2, 3, 4). Furthermore, aberrations in signaling pathways regulating cell migration contribute to tumor invasion (5) and metastasis (6). Over the last decades, two main modes of migration have been identified: adhesion-dependent mesenchymal (7) and adhesion-independent amoeboid migration (8). These migration modes differ in the way forces are generated and transduced within the cell. Importantly however, the breaking of cell symmetry is a fundamental process at the basis of any migration event (9, 10).

In the absence of external polarity cues, several mechanisms of spontaneous symmetry breaking have been proposed and are based on polarization of cytoskeleton components (11). For instance, gradients or patterns of morphogens can arise due to specific reaction-diffusion patterns within the cell, leading to its polarization (12). More recently, several mechanisms of spontaneous symmetry breaking of the actin myosin system itself have been proposed, based either on actin polymerization (13, 14) or acto-myosin contractility (15, 16, 17). However, relating such symmetry breaking events of the various components of the cellular cytoskeleton to both cell-substrate forces and cell locomotion remains largely unexplored.

In the specific case of mesenchymal migration, the spatio-temporal sequence of mechanical symmetry breaking remains controversial. Different models are distinguished by the temporal order in which distinct cytoskeleton forces are activated to trigger directional movement (18). Most studies emphasize force generation due to actin polymerization in the cell front as a first step to initiate migration (3, 19). On the contrary, acto-myosin II-mediated contractility within the cell rear has been identified as a first step to break cell symmetry in keratocytes (20). Thus, determining the spatio-temporal dynamics of cellular forces and morphological events at the initiation of a migration is still an open and major question in biology.

To investigate quantitatively the dynamics of spontaneous symmetry breaking events in cells at the level of both morphological parameters and distribution of interaction forces with the environment, we developed a one-dimensional migration assay (**Fig. 1A**) that combined time-resolved traction force microscopy (TFM) (21, 22, 23) and soft micro patterning (24) (Materials and methods can be found in supplementary materials).

**Fig. 1.**
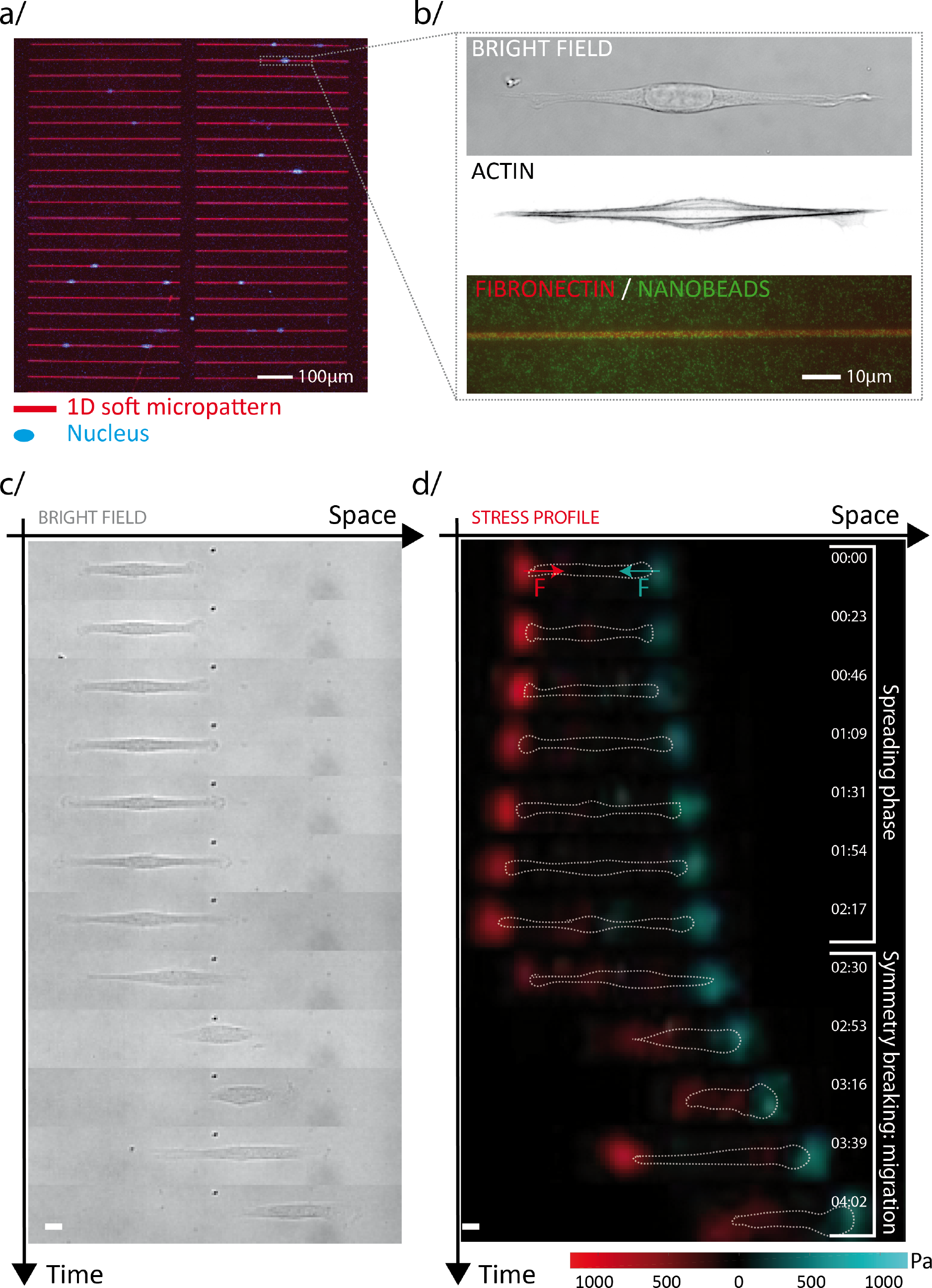
One-dimensional single cell migration assay based on soft micropatterning and traction force microscopy mimics complex 3D fibrillar *in vivo* migration. **(a**) 40 kPa polyacrylamide gel with RPE1 cells (blue: nucleus staining) on top of 2 *μ*m micro-patterned fibronectin lines (red). **(b**) Brightfield, actin cytoskeleton and bead imaging of RPE1 on 2 *μ*m line allowed extracting morphometric and mechanical parameters simultaneously. **(c**) Time sequence of RPE1 cell migrating on fibronectin lines and **(d**) its associated stress profile extracted via TFM (dotted white line: cell outline; color coded stress profile depending on the direction of applied traction forces 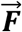: red in and blue against the direction of migration; Scale bars: 10 μm).

Using this bottom-up approach, we followed single epithelial cells (hTERT-immortalized retinal pigment epithelial cell line [RPE1]) during the initiation of spontaneous migration and extracted morphometric and mechanical parameters. As expected (25, 26), RPE1 cells plated on patterned 40kPa polyacrylamide hydrogels adhered to one-dimensional fibronectin lines (2 or 5 *μ*m width) within 1-2 hours. The cells displayed elongated shapes with long actin fibers oriented parallel to the micropattern and cell axis (**Fig. 1B**).

In the absence of any external cue, we observed a biphasic motile behavior: symmetric elongation of a static cell (spreading phase) prior to spontaneously initiated directional movement (migration phase) (**Fig. 1C**). In parallel, tangential stress measurements revealed defined stress compartments at both cell edges due to contractile forces oriented towards the center of the cells (**Fig. 1D**). Hence, cells behaved as force dipoles, as described previously (14, 27, 28, 29). During the spreading phase, both cell elongation dynamics and force distribution patterns were fully symmetric with respect to the cell center of mass. At the onset of motility, morphological polarization and simultaneous asymmetrical redistribution of forces occurred, characterized by a single defined local stress compartment at the cell front and a significantly widened stress distribution with lower traction stress at the rear (**Fig. 1D**). This was accompanied by rapid retraction of the cell rear (**Fig. 1D**).

Current models emphasize the formation of a distinct cell front as the first event when cell migration is initiated (10, 30). In contrast, we observed that cell spreading was qualitatively symmetric on both sides and that symmetry breaking occurred with the sudden retraction of the rear. This led us to hypothesize that contractility builds up in a non-polarized cell, resulting in a local stress increase at both extremities.

To challenge the hypothesis that symmetry breaking does not require pre-established rear-front polarity as previously thought (31, 20), we quantified the coordination between mechanical polarization and morphological events. To first confirm the qualitative observation of anisotropic redistribution of traction forces, we adapted multipole analyses, classically used in the field of micro-swimmers (32), to quantify the asymmetry of the force distribution. We first projected the stress profile along the micropattern axis to obtain a 1D stress profile, a mechanical footprint of the cell. From that, we computed the variance of (+)- and (-)-directed traction stress profiles (D+, D-), which quantified the spatial distribution of each stress compartment at opposite poles of the cell. The normalized ratio, (D+ - D-)/(D+ + D-), (analogous to the normalized quadrupole) quantifies the symmetry of the spatial stress distribution and will be referred to as force asymmetry parameter (**Fig. 2A**).

**Fig. 2.**
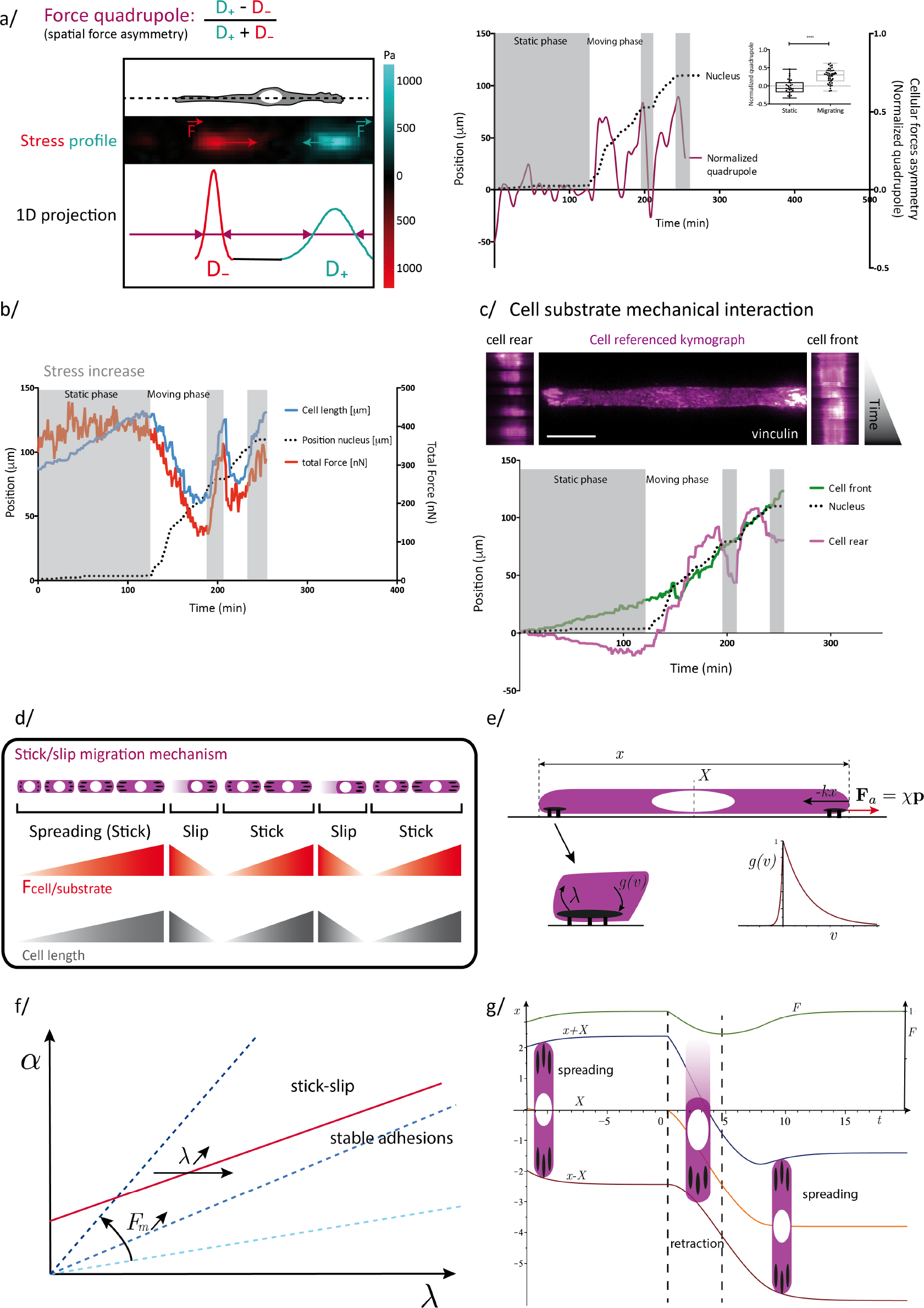
RPE1 cells exhibit intermittent migration following a stick-slip motion. **(a**) Scheme of the force asymmetry analysis: the normalized quadrupole was extracted from the 1D projection of the stress profile of an adherent cell (color coded stress map and 1D profile depending on the direction of applied traction forces 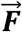 exerted: red in and blue against the direction of migration). Dynamic measurements revealed a symmetric spatial force profile during static spreading and an asymmetric distribution during migration phases. Inset: average force asymmetry during static and mobile phases of several cells (n = 10). **(b**) Cell length and total force correlation: increase during spreading phase and decrease during migration. **(c**) Referenced kymograph of RPE1 cells stably expressing vinculin-eGFP showing a continuous attachment of the front while adhesions in the rear detached and reattached during one migration cycle (Scale bar: 10 μm). Tracking the front, rear and nucleus position over time could further represent this destabilization of the rear. **(d**) Deduced scheme of the proposed stick-slip migration mechanism: during non-motile spreading (stick) the cell builds up a high traction force that eventually will overcome adhesion strength in the perspective rear of the cell. Upon the retraction of the rear, the cell shortens and lowers its mechanical interaction with the substrate to initiate migration (slip). **e,** Schematic of the model and parameters as defined in the text. **(f**) Example of stick slip dynamics predicted by the model. Dynamical equations 2-3 are solved numerically with *vm* = *0.5*, ***ν*_*p*_**= *0.5*, *λ* = *1*, ***μ*** = *1*, *α* = *1* (arbitrary units). Blue, orange and brown line show rear, nucleus and front position over time, respectively. Green line depicts the relative traction force level F. **(g**) Phase diagram of dynamic behaviors predicted by the model, as a function of the actin turn-over rate *λ* and phenomenological parameter *α* (arbitrary units). Dashed-lines show different values of the maximal contractile force 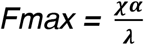.

Non-migrating cells exhibited a force asymmetry parameter fluctuating around zero, indicating a non-polarized static phase (**Fig. 2A**). Consistently, fluctuations in the actin profiles were also observed in static phases (**Fig. S1**). Importantly, no significant polarization of actin distribution was observed prior to migration initiation. Nevertheless, upon initiation of each migration step, the force asymmetry parameter displayed a sharp transient peak. This sudden increase corresponded to a widening of the spatial stress distribution in the rear of the cell while the stress pattern at the cell front remained localized to the cell edge. This asymmetry subsequently relaxed leading to another static phase. Several iterations of such phases were typically observed. Consistently, we found larger values in the amplitude of the asymmetry parameter in moving phases in comparison to the static ones for multiple analyzed cells. Thus, initiation of migration is characterized by a sharp increase of the force asymmetry parameter and can occur in absence of prior polarization of the actin cytoskeleton.

We subsequently hypothesized that stress builds up and fluctuates during the spreading phase until one end randomly detaches producing a cell rear. This hypothesis was supported by the evolution of the total traction force, a measure of the strength of the mechanical interaction of the cell with the substrate, quantified via TFM. We observed that, in static phases, cell spreading was associated with an increase of the total traction force. Upon the initiation of migration, the force level dropped approximately by 50% (**Fig. S2**). Strikingly, this decrease in mechanical interaction was directly correlated with a shortening in cell length due to the sudden retraction of the rear (**Fig. 2B**, **Fig. S3**). To confirm the role of adhesion detachment, we fluorescently labeled cell-substrate anchor points using vinculin-eGFP to follow the time evolution of adhesion patches during migration. Adhesion sites at the front of the cell were continuously contacting the substrate while adhesion sites at the rear followed two distinct phases: attachment (cluster growth) and switching abruptly to detachment (disassembly and sliding of smaller adhesion patches) (Fig. 2C). Cell morphology and its polarity features showed similar behavior as after the initial symmetric spreading phase, abrupt retraction of the rear triggered subsequent nuclear translocation. Furthermore, throughout the migration cycle the trailing edge displayed two distinct phases of motion, while the front continuously moved forward (**Fig. 2C**). This destabilization of the trailing edge demonstrated the critical role of adhesion detachment in the back of the cell. The observed discontinuous migration is known in physics as a stick-slip mechanism (**Fig. 2D).** During the initial spreading phase, cells elongated symmetrically while increasing their contractile stress (stick). Upon reaching a level of stress that adhesion complexes could no longer sustain, adhesions on one cell edge stochastically detached from the substrate. This led to cell shortening due to retraction of the rear and a decrease in cell-substrate interaction (slip). Recovery of the initial cell length and contractility level occurred during the subsequent stick phase. As a consequence of this stick slip migration, the propensity of cells to enter migratory phases appeared to crucially depend on (i) contractility and (ii) adhesion properties.

In order to substantiate this observed stochastic stick-slip behavior, we devised a physical model based on minimal ingredients. The actin cytoskeleton was described as an active, homogeneous 1D viscoelastic gel (33). We assumed that the cell’s cytoskeleton was fully unpolarized, and that the cell body could be mechanically characterized by an effective stiffness *k*. This elastic behavior encompasses active (i.e. due to motor activity) and passive contributions of both cytoskeleton and membrane. Adhesion sites were described in the framework of the active gel theory as localized regions at both cell extremities carrying outward pointing actin polarity ***p***, and subjected to an active force ***F***_***a***_=*χ* ***p***, where *χ* is a phenomenological coupling constant, which induced cell expansion. The key ingredient of the model relies on the dynamics of adhesion sites, which was written phenomenologically as *ṗ* = *g*(*v*_*p*_) − *λp*. Here *λ* models the rate of actin turnover, and *g* the dynamics of adhesion sites assembly that depends on the local velocity *v*_*p*_ = *v* · *u*_*p*_ over the substrate. Importantly, *g* is a priori very asymmetric (**Fig. 2E**). This accounts for the fact that adhesion assembly is drastically reduced upon edge retraction, and mildly affected by edge expansion. The analysis of the model revealed that the actin turnover rate critically controls the dynamics. In particular at slow turnover rate (as defined in SI), the system was found to display a stochastic stick-slip behavior, (which notably differs from classical stick-slip behaviors characterized by deterministic oscillations). Cells were predicted to slowly expand and reach the fixed point of the dynamics where any fluctuation leading to infinitesimal retraction is unstable: one end of the cell therefore retracts before spreading symmetrically again. Finally, the model successfully predicts that it is critically controlled by adhesion turnover rate *λ* and maximal contractile force, as summarized in the phase diagram of **Fig. 2F**, and reproduces the observed stochastic stick-slip dynamics (**Fig. 2G**).

To challenge the proposed stochastic stick-slip mechanism, we used optogenetics to disrupt its predicted spatio-temporal sequence. We used NIH3T3 cells stably expressing a Cry2-CIBN optogenetic probe to dynamically control the localization of ArhGEF11, an upstream regulator of the master regulator of cell rear retraction, RhoA (from now on referred to as optoGEF-RhoA) (34). Upon stimulation with blue light, optoGEF-RhoA dimerizes with the CAAX-anchored protein CIBN, leading to its immediate translocation from the cytoplasm to the membrane where it activates RhoA, triggering asymmetric recruitment of actin and subsequent cell migration away from the photo-activation spot. The initiated movement was characterized by a distinct front-rear polarity that was maintained throughout the whole stimulation cycle. Interestingly, by switching the side of stimulation, actin polarity and direction of movement were inverted (**Fig. 3A**).

**Fig. 3.**
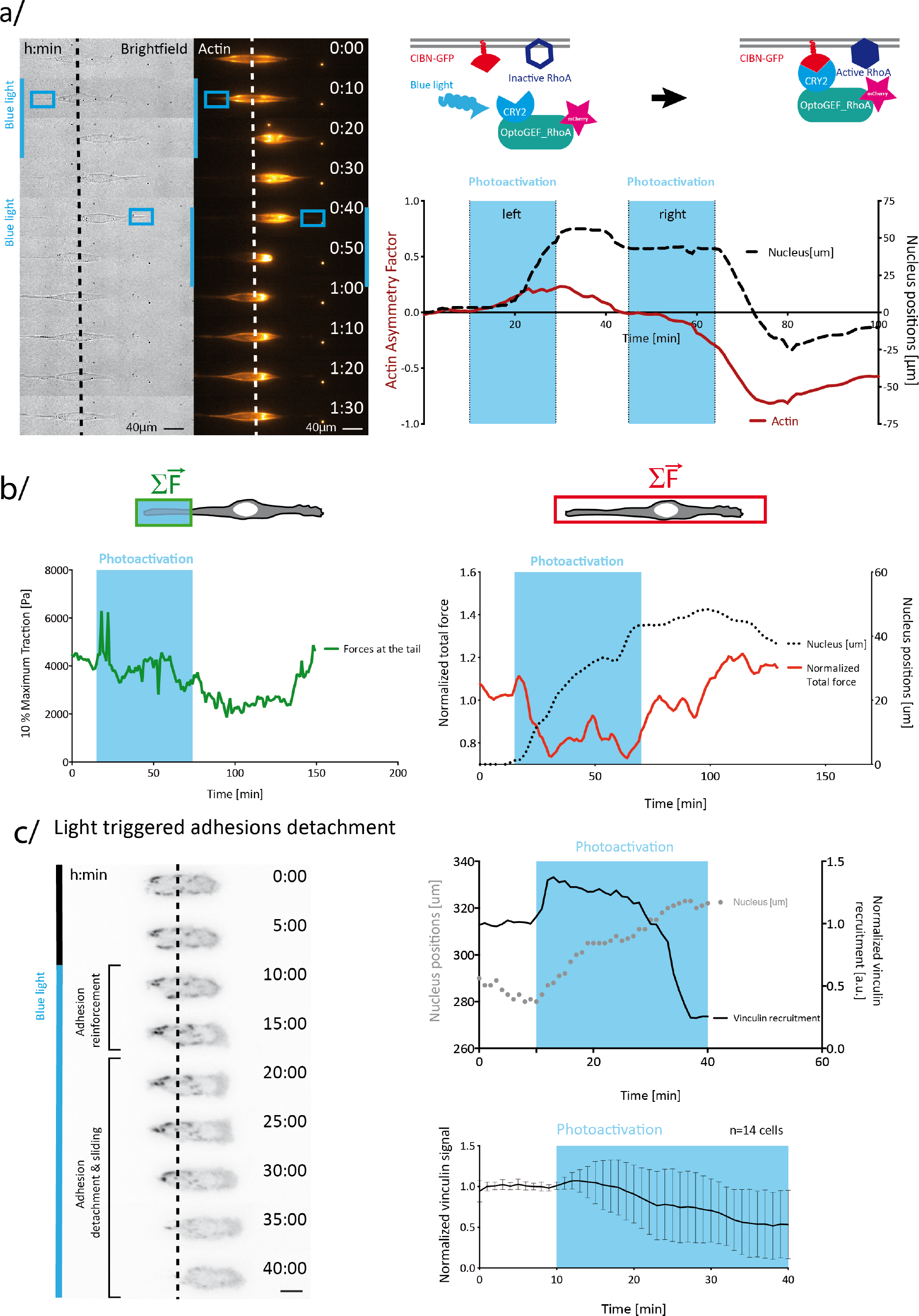
RhoA optogenetic control of cellular force symmetry breaking. **(a**) Schematic representation of light-induced Cry2-CIBN dimerization and local RhoA activation due to its close proximity to its upstream regulator opto_GEFRhoA. Brightfield and actin imaging and quantification showed the light induced migration away from the photoactivation area (blue square), which is characterized by a transient front-rear polarity and actin asymmetry (dashed line: nucleus position at t0). **(b**) Local and global force response of the light-induced rear and of the whole cell, respectively, showed a transient local contractility increase at the perspective rear followed by a global decrease of the mechanical cell-substrate interaction. **(c**) stably expressing vinculin-iRFP revealed local adhesion reinforcement within the photostimulated area followed by a subsequent adhesion detachment. Dashed line indicates nucleus position at t0. Scale bar: 10 μm.

This optogenetic approach combined with quantitative force measurements revealed a RhoA-mediated instantaneous and local increase of traction forces in the zone of activation. This transient and spatially confined force increase was followed by a global decrease of the mechanical interaction of the moving cell with its substrate, as seen on the total traction force (**Fig. 3B**). This drop was similar to the one observed during spontaneous migration (**Fig. 2C**), which was attributed to adhesion detachment at the cell rear. To confirm that the same process was at play here we imaged adhesions by transiently transfecting vin-iRFP. Upon light-induced RhoA activation, we observed first reinforcement, then detachment and sliding of adhesions (**Fig. 3C**). Indeed, as acto-myosin contractility was stimulated, adhesions were submitted to an increasing level of stress that first led to vinculin recruitment (positive feedback) (35), but ultimately caused the adhesion to dissociate. Hence, local stimulation artificially created the cell rear, triggering the first steps of cell translocation (adhesion detachment) as in the case of spontaneous migration.

A key prediction of the stick-slip model is that spontaneous symmetry breaking strongly depends on contractility and adhesiveness. To challenge this prediction and to further investigate the stick-slip migration mechanism illustrated in **Fig. 2**, we systematically analyzed the main parameters of our theoretical model (cell length, adhesion size, and total traction forces) and correlated them with the migratory behavior of single cells of two distinct cell types exhibiting different migratory behavior. The instantaneous speed of the cell centroid averaged over the whole cell trajectory was used as a parameter to represent the migration capacity of single cells. To test the broader applicability of the model, fast-migrating RPE1 (36) cells were compared to fibroblast cells (NIH3T3) that exhibit slow mesenchymal migration (37).

RPE1 cells exhibited a higher speed compared to NIH3T3 that mostly remained in a static spreading phase with less frequent retraction phases. Comparing cell morphology and traction force level of both cell types, we observed that NIH3T3 cells exhibited a longer spreading length associated with a larger mechanical interaction of the cells with their microenvironment (**Fig. 4A**). This result may appear counter-intuitive as larger traction forces should facilitate detachment of adhesions and thus cellular movement. However, in the classical catch-bond model, an increase of force would also induce a stabilization and reinforcement of adhesion sites (38). Consistent with this, NIH3T3 cells had larger adhesion patches compared to RPE1 cells.

**Fig. 4.**
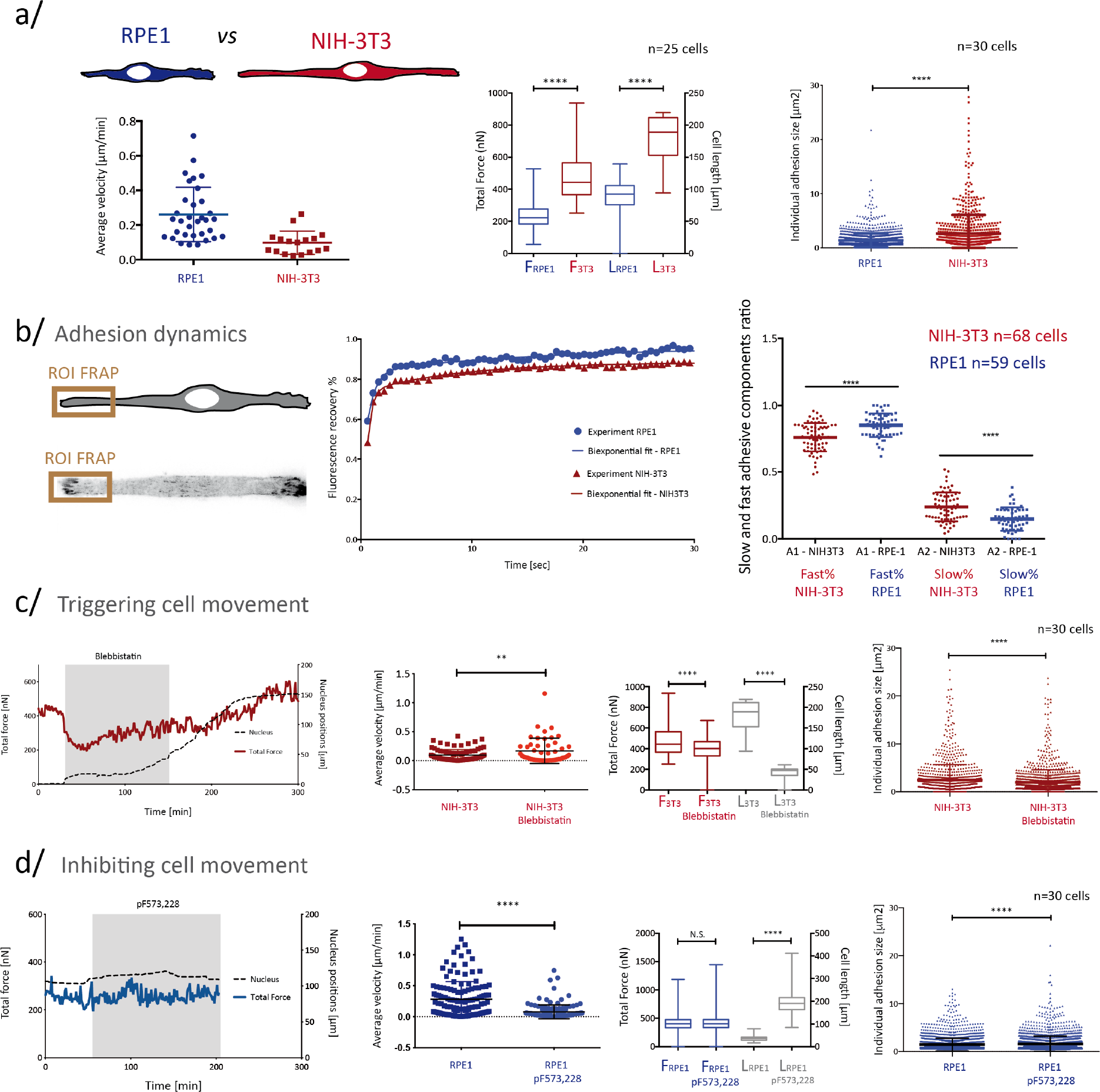
Adhesiveness and contractility control the migratory behavior of NIH3T3 and RPE1 cells. **(a**) Comparison of instantaneous migration speed, total force, cell length and individual adhesion size of RPE1 and NIH3T3 cells. **(b**) FRAP experiments of adhesions located at one cell edge were modeled with a bi-exponential fit to extract a fast and slow component representing mobile vinculin within the cytoplasm and slow vinculin bound to adhesions. **(c) and (d**) Altering the migratory behavior of RPE1 and NIH3T3 using 1 *μ*M pF573,228 to inhibit and 3 *μ*M blebbistatin to trigger migration, respectively. Shown are measured parameters relevant for stick-slip migration: average migration speed, total force, cell length and individual adhesion size. Statistical significance tested with unpaired t-test (*P* < 0.05). Scatter plots with mean and standard deviation. Box plots from minimum and maximum values with the mean and standard deviation. Number *n* of analyzed cells per condition indicated on the respective graph figures.

To analyze adhesion strength in more detail, we quantified adhesion dynamics in both cell types. First, total internal reflection fluorescence microscopy (TIRF) of vin-eGFP adhesions revealed faster adhesion turnover in RPE1 cells compared to NIH3T3 fibroblasts (**Movie S1**). Fluorescence recovery after photobleaching (FRAP) experiments over single adhesion patches localized at the cell edges of each cell type revealed two time components: a fast one that was related to the diffusion of vinculin molecules within the cytosol and a slow one corresponding to the residence time of immobilized vinculin within the adhesion sites (**Fig. 4B**). The measured slow and fast component ratios revealed that RPE1 cells displayed a lower fraction of bound vinculin compared to NIH3T3. Since vinculin binding promotes adhesion stability, our data indicated that RPE1 cells exhibited more labile adhesions, while NIH3T3 adhesions were expected to sustain higher tension without breaking. These findings are in agreement with the stick-slip model since faster RPE1 cells would undergo fast spreading/retraction cycles (large *λ*), while less motile NIH3T3 relaxed more slowly to the unstable fixed point (small *λ*). Therefore, the migratory behavior of these two cell types could be explained, in the framework of a stick-slip model, by cells having different levels of adhesiveness and contractility.

To further confirm the validity of this model, we used pharmacological treatments to perturb the balance between adhesiveness and contractility. We used a low dose of blebbistatin (3μM) to decrease contractility (39) in NIH3T3 fibroblasts and 1μM pF573,228 to stabilize adhesions (40) in RPE1 cells. As both parameters (contractility and adhesion strength) are bi-directionally coupled through positive feedback loops (38, 41, 42), one could not be modulated without affecting the other. Blebbistatin-treated NIH3T3 cells were able to initiate migration more readily, as shown by the increase of their migration speed (**Fig. 4C**). They exhibited a decrease of total traction force as expected, but also a shortening of the average cell length, which suggested that these cells can more easily detach their adhesions. Indeed, the size of adhesion patches decreased significantly upon blebbistatin treatment (**Fig. 4C**). Hence, by inhibiting contractility, cell adhesiveness was lowered, which facilitated the rear detachment and led to cell shortening and increased motility. In agreement with the stick-slip model, low maximal contractile force corresponded with low cell/substrate interactions, giving rise to reduced cell spreading and therefore smaller cell length and potentially larger speeds, provided that the cytoskeleton is polarized.

On the contrary, stabilizing focal adhesions on RPE1 cells decreased their average velocity. It also induced a lengthening of the cells and larger adhesion patches (**Fig. 4D**) as predicted by our model: diminishing the turnover rate *λ* induces a marked stick-slip behavior, with long spreading phases, and therefore large cell length, and slow speed. Remarkably, the dependence of the stick-slip behavior on the turnover rate and contractility results in inverse correlation between average cell length and migration velocity (**Fig. 5A**), which was consistently observed in both NIH3T3 and RPE1 cells. More elongated cells, such as NIH3T3, were associated with stronger adhesions, as they could spread more without detaching, and hence a lower velocity. When this detachment occurred at an early stage of spreading, corresponding to low stress levels, cells were shorter and exhibited higher migration speeds, as in the case of RPE1.

**Fig. 5.**
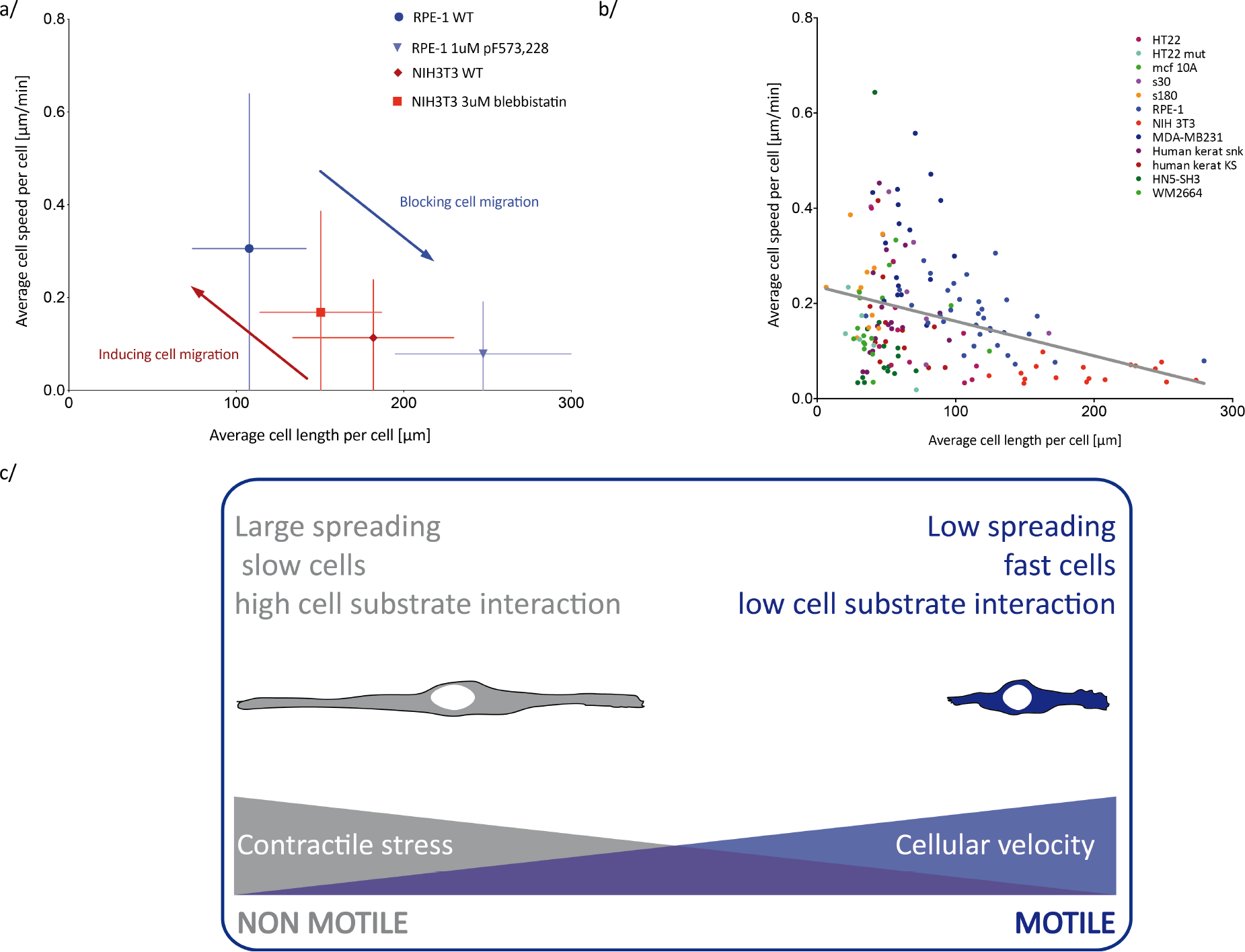
The inverse relation between cell length and speed. **(a**) Experimentally deduced phase diagram using a pharmacological approach to alter the migratory behavior of RPE1 and NIH3T3 cell (error bars show the standard deviation from the mean). **(b**) Length-speed relation validated by analyzing several cell types coming from the cell race data (one color used per cell type; black line: linear fit of all data points). **(c**) Summary showing how cell contractility, and therefore adhesiveness and cell length, control cellular migration.

Finally, we asked if the stick-slip paradigm would operate also in the presence of other polarization mechanisms. We utilized deposited data of single cell trajectories of various cell types on patterned adhesive one-dimensional lines (World Cell Race (37)). For each cell line, both instantaneous cell speed and cell length were extracted and correlated with each other (**Fig. 5B**). Strikingly, the negative correlation between cell length and cell speed, consistent with the stick-slip regime, was confirmed for most of cell lines.

Our findings demonstrate that a stochastic stick-slip mechanism, which is intrinsically based on the properties of adhesion dynamics, is a very robust feature of adherent cell migration. In particular, while this mechanism provides a simple scenario of spontaneous symmetry breaking and cell polarization, our results suggest that stick-slip behavior occurs also in the presence of other polarization mechanisms.

Using a one-dimensional approach based on soft micropatterning, force imaging and optogenetics in combination with theoretical approaches, we have uncovered a generic, stick-slip mechanism that can initiate cell migration. This mechanism allows cells to spontaneously break their symmetry by stochastically detaching adhesive contacts on one side, resulting in a migratory step in the opposite direction. This work shows that cell symmetry breaking can emerge independently of a prior polarity of the actin cytoskeleton, due to instabilities of the mechanochemical coupling of the cell to its environment via adhesion sites. This process is found to be controlled by the interplay of contractile forces and focal adhesion dynamics. Hence, by modifying contractility and adhesiveness of the cell, the rate of such stochastic steps (i.e. the instantaneous speed of cell motion) can be controlled. Interestingly, we found that stochastic stick slip is responsible for a negative correlation between cell length and cell speed, which we observed across many cell types, thereby further emphasizing the relevance and robustness of this mechanism. In the light of our findings, cell length represents a direct readout of cell adhesiveness and thus appears as straightforward parameter to predict cell migratory behavior.

We observed for several cells of different cell types that the first stochastic step can lead to the emergence of a more persistent polarity within the moving cell, as some cells tend to take several migration steps in the same direction. Stochastic stick slip therefore appears as a basic mechanism of symmetry breaking conserved across adherent mammalian cell types, which can coexist with other polarization mechanisms, based e.g. on cytoskeleton instabilities or reaction diffusion patterns.

## Acknowledgements

We thank Matthieu Piel and Paolo Maiuri for valuable discussions and sharing the cell race data with us. We also thank Thomas Boudou and Manuel Théry for critical discussions, Jean Bernard for technical assistance, and members of the MOTIV team at LiPhy for support. We thank Laurent Blanchoin’s Cytomorpholab in Grenoble for providing us RPE1 cells and Helder Maiato from the University of Porto for providing us with NIH3T3 cells. Furthermore, we want to thank Alexander Kyumurkov at the Institute of Advanced Biology in Grenoble for assisting with adhesion imaging.

## Funding

This work was supported by Nanoscience fondation (MB), and the ARC fondation (MB).

## Author contributions

K.H. performed experiments and analyzed the data. S.dB. performed optogenetic experiments on vinculin-iRFP transfected cells. L.V. designed the optogenetic cell line in M.C.’s laboratory. Y.A.M. designed the vinculin-eGFP RPE1 and NIH3T3 cells in C.A.R.’s laboratory. C.F. provided insights on the FRAP experiments and performed the related data analysis. R.V. developed the theoretical framework. M.B. supervised the research. All authors contributed to write the paper.

## Competing interests

No authors have competing interests.

## List of Supplementary Materials

Materials and Methods Figure S1 - S3

Movie S1

References 24, 27, 43 - 46

## References

1. Renkawitz, J. and Sixt, M., Mechanisms of force generation and force transmission during interstitial leukocyte migration. EMBO reports 11 (10), 744–750 (2010).

2. Webb, D.J., Parsons, J.T. and Horwitz, A.F., Adhesion assembly, disassembly and turnover in migrating cells–over and over and over again. Nature Cell Biology 4 (4), E97 (2002).

3. Ridley, A.J., Schwartz, M.A., Burridge, K., Firtel, R.A., Ginsberg, M.H., Borisy, G., Parsons, J.T. and Horwitz, A.R., Cell migration: integrating signals from front to back. 302 (5651), 1704–1709 (2003).

4. Friedl, P. and Wolf, K., Plasticity of cell migration: a multiscale tuning model. The Journal of cell biology, jcb-200909003 (2009).

5. Monzo, P., Chong, Y.K., Guetta-Terrier, C., Krishnasamy, A., Sathe, S.R., Yim, E.K., Ng, W.H., Ang, B.T., Tang, C., Ladoux, B. and Gauthier, N.C., Mechanical confinement triggers glioma linear migration dependent on formin FHOD3. Molecular biology of the cell 27 (8), 1246–1261 (2016).

6. van Zijl, F., Krupitza, G. and Mikulits, W., Initial steps of metastasis: Cell invasion and endothelial transmigration. Mutation Research/Reviews in Mutation Research 728 (1), 23–34 (2011).

7. Huttenlocher, A. and Horwitz, A.R., Integrins in cell migration. Cold Spring Harbor perspectives in biology 3 (9), a005074 (2011).

8. Fackler, O.T. and Grosse, R., Cell motility through plasma membrane blebbing. The Journal of cell biology 181 (6), 879–884 (2008).

9. Lomakin, A. J., Lee, K. C., Han, S. J., Bui, D. A., Davidson, M., Mogilner, A., & Danuser, G., Competition for actin between two distinct F-actin networks defines a bistable switch for cell polarization. Nature Cell Biology 17 (11), 1435 (2015).

10. Lauffenburger, D. A., & Horwitz, A. F., Cell migration: a physically integrated molecular process. Cell 84 (3), 359–369 (1996).

11. Tee, Y.H., Shemesh, T., Thiagarajan, V., Hariadi, R.F., Anderson, K.L., Page, C., Volkmann, N., Hanein, D., Sivaramakrishnan, S., Kozlov, M.M. and Bershadsky, A.D., Cellular chirality arising from the self-organization of the actin cytoskeleton. Nature cell biology 17 (4), 445 (2015).

12. Turing, A. M., The chemical basis of morphogenesis. Philosophical Transactions of the Royal Society of London 237 (641), 37–72 (1952).

13. Callan Jones, A.C., Joanny, J.F. and Prost, J., Viscous-fingering-like instability of cell fragments. Physical review letters 100 (25), 258106 (2008).

14. Blanch-Mercader, C. and Casademunt, J., Spontaneous Motility of Actin Lamellar Fragments. Physical review letters 110 (7), 078102 (2013).

15. Hawkins, R.J., Piel, M., Faure-Andre, G., Lennon-Dumenil, A.M., Joanny, J.F., Prost, J. and Voituriez, R., Pushing off the walls: a mechanism of cell motility in confinement. Physical review letters 102 (5), 058103 (2009).

16. Callan Jones, A.C. and Voituriez, R., Active gel model of amoeboid cell motility. New Journal of Physics 15 (2), 025022 (2013).

17. Ruprecht, V., Wieser, S., Callan-Jones, A., Smutny, M., Morita, H., Sako, K., Barone, V., Ritsch-Marte, M., Sixt, M., Voituriez, R. and Heisenberg, C.P., Cortical contractility triggers a stochastic switch to fast amoeboid cell motility. Cell 160 (4), 673–685 (2015).

18. Cramer, L. P., Forming the cell rear first: breaking cell symmetry to trigger directed cell migration. Nature Cell Biology 12 (7), 628 (2010).

19. Pollard, T.D. and Borisy, G.G., Cellular Motility Driven by Assembly and Disassembly of Actin Filaments. Cell 112 (4), 453–465 (2003).

20. Yam, P.T., Wilson, C.A., Ji, L., Hebert, B., Barnhart, E.L., Dye, N.A., Wiseman, P.W., Danuser, G. and Theriot, J.A., Actin–myosin network reorganization breaks symmetry at the cell rear to spontaneously initiate polarized cell motility. Journal of Cell Biology 178 (7), 1207–1221 (2007).

21. Butler, J.P., Tolic-Nørrelykke, I.M., Fabry, B. and Fredberg, J.J., Traction fields, moments, and strain energy that cells exert on their surroundings. American Journal of Physiology-Cell Physiology 282 (3), C595–C605 (2002).

22. Dembo, M. and Wang, Y.L., Stresses at the cell-to-substrate interface during locomotion of fibroblasts. Biophysical journal 76 (4), 2307–2316 (1999).

23. Lafaurie-Janvore, J., Maiuri, P., Wang, I., Pinot, M., Manneville, J.B., Betz, T., Balland, M. and Piel, M., ESCRT-III assembly and cytokinetic abscission are induced by tension release in the intercellular bridge. Science 339 (6127), 1625–1629 (2013).

24. Tseng, Q., Wang, I., Duchemin-Pelletier, E., Azioune, A., Carpi, N., Gao, J., Filhol, O., Piel, M., Théry, M. and Balland, M., A new micropatterning method of soft substrates reveals that different tumorigenic signals can promote or reduce cell contraction levels. Lab on a Chip 11 (13), 2231–2240 (2011).

25. Doyle, A. D., Wang, F. W., Matsumoto, K., & Yamada, K. M., One-dimensional topography underlies three-dimensional fibrillar cell migration. The Journal of cell biology 184 (4), 481–490 (2009).

26. Schuster, S.L., Segerer, F.J., Gegenfurtner, F.A., Kick, K., Schreiber, C., Albert, M., Vollmar, A.M., Rädler, J.O. and Zahler, S., Contractility as a global regulator of cellular morphology, velocity, and directionality in low-adhesive fibrillary micro-environments. Biomaterials 102, 137–147 (2016).

27. Mandal, K., Wang, I., Vitiello, E., Orellana, L.A.C. and Balland, M., Cell dipole behaviour revealed by ECM sub-cellular geometry. Nature communications 5, 5749 (2014).

28. Stéphanou, A., Le Floc’h, S. and Chauvière, A., A hybrid model to test the importance of mechanical cues driving cell migration in angiogenesis. Mathematical Modelling of Natural Phenomena 10 (1), 142–166 (2015).

29. Bergert, M., Erzberger, A., Desai, R.A., Aspalter, I.M., Oates, A.C., Charras, G., Salbreux, G. and Paluch, E.K., Force transmission during adhesion-independent migration. Nature cell biology 17 (4), 524–529 (2015).

30. Mitchison, T.J. and Cramer, L.P., Actin-based cell motility and cell locomotion. Cell 84 (3), 371–379 (1996).

31. Ladoux, B., Mège, R.M. and Trepat, X., Front–rear polarization by mechanical cues: From single cells to tissues. Trends in Cell Biology 26 (6), 420–433 (2016).

32. Wu, H., Thiébaud, M., Hu, W.F., Farutin, A., Rafaï, S., Lai, M.C., Peyla, P. and Misbah, C., Amoeboid motion in confined geometry. Physical Review E 92 (5), 050701 (2015).

33. Kruse, K., Joanny, J.F., Jülicher, F., Prost, J. and Sekimoto, K., Generic theory of active polar gels: a paradigm for cytoskeletal dynamics. The European Physical Journal E 16 (1), 5–16 (2005).

34. Valon, L., Etoc, F., Remorino, A., di Pietro, F., Morin, X., Dahan, M. and Coppey, M., Predictive spatiotemporal manipulation of signaling perturbations using optogenetics. Biophysical journal 109 (9), 1785–1797 (2015).

35. Galbraith, C.G., Yamada, K.M. and Sheetz, M.P., The relationship between force and focal complex development. Journal of Cell Biology 159 (4), 695–705 (2002).

36. Maiuri, P., Rupprecht, J.F., Wieser, S., Ruprecht, V., Bénichou, O., Carpi, N., Coppey, M., De Beco, S., Gov, N., Heisenberg, C.P. and Crespo, C.L., Actin flows mediate a universal coupling between cell speed and cell persistence. Cell 161 (2), 374–386 (2015).

37. Maiuri, P., Terriac, E., Paul-Gilloteaux, P., Vignaud, T., McNally, K., Onuffer, J., Thorn, K., Nguyen, P.A., Georgoulia, N., Soong, D. and Jayo, A., The first world cell race. 22 (17) (2012).

38. Liu, Z., Bun, P., Audugé, N., Coppey-Moisan, M. and Borghi, N., Vinculin head– tail interaction defines multiple early mechanisms for cell substrate rigidity sensing. Integrative Biology 8 (6), 693–703 (2016).

39. Kovács, M., Tóth, J., Hetényi, C., Málnási-Csizmadia, A. and Sellers, J.R., Mechanism of Blebbistatin Inhibition of Myosin II. Journal of Biological Chemistry 279 (34), 35557–35563 (2004).

40. Slack-Davis, J.K., Martin, K.H., Tilghman, R.W., Iwanicki, M., Ung, E.J., Autry, C., Luzzio, M.J., Cooper, B., Kath, J.C., Roberts, W.G. and Parsons, J.T., Cellular characterization of a novel focal adhesion kinase inhibitor. Journal of Biological Chemistry 282 (20), 14845–14852 (2007).

41. Renkawitz, J., Schumann, K., Weber, M., Lämmermann, T., Pflicke, H., Piel, M., Polleux, J., Spatz, J.P. and Sixt, M., Adaptive force transmission in amoeboid cell migration. Nature cell biology 11 (12), 1438 (2009).

42. Liu, Y.J., Le Berre, M., Lautenschlaeger, F., Maiuri, P., Callan-Jones, A., Heuzé, M., Takaki, T., Voituriez, R. and Piel, M., Confinement and low adhesion induce fast amoeboid migration of slow mesenchymal cells. Cell 160 (4), 659–672 (2015).

43. Tse, J.R. and Engler, A.J., Preparation of hydrogel substrates with tunable mechanical properties. Current protocols in cell biology, 10–16 (2010).

44. Sabass, B., Gardel, M.L., Waterman, C.M. and Schwarz, U.S., High Resolution Traction Force Microscopy Based on Experimental and Computational Advances. Biophysical journal 94 (1), 207–220 (2008).

45. Vignaud, T., Ennomani, H. and Théry, M., Polyacrylamide hydrogel micropatterning‥ Methods in cell biology 120, 93–116 (2014).

46. Horzum, U., Ozdil, B. and Pesen-Okvur, D., Step-by-step quantitative analysis of focal adhesions. MethodsX 1, 56–59 (2014).

